# Multiparametric *in vivo* mapping reveals tissue-specific mitochondrial aging trajectories

**DOI:** 10.64898/2026.09.09.750228

**Authors:** Wenyu Tang, Haoyu Xu, Steven Beck, Marcus T Cicerone, Sung Min Han, Wei-Wen Chen

## Abstract

Mitochondrial dysfunction is a hallmark of aging, yet how mitochondrial states are remodeled across tissues and subcellular compartments *in vivo* remains elusive. Progress has been limited, in part, because mitochondrial physiology is highly sensitive to experimental perturbations, underscoring the need for minimally disruptive measurement strategies. Here, we establish a tissue-resolved, *in vivo* framework for the quantitative analysis of mitochondrial states in live, intact *Caenorhabditis elegans* without confounding effects from mounting-induced hypoxia. This platform couples two-photon fluorescence lifetime imaging microscopy (2p-FLIM) with a custom segmentation pipeline, MitoSLIT, to track functional and structural features across multiple tissues and single neurons. By integrating membrane potential-associated TMRM intensity, lifetime-based microenvironmental metrics, and morphological descriptors, we uncover localized metabolic heterogeneity masked by conventional intensity analysis. Leveraging this framework, we mapped physiological aging against mitochondrial shifts induced by acute stress and fission-fusion mutations. Our analyses reveal that mitochondrial aging is highly tissue-specific, executing distinct trajectories across cell types. Extending the framework to genetically identified neurons revealed age-dependent divergence between somatic and axonal mitochondrial states, accompanied by structural remodeling and a late shift in optical redox ratio. Together, our findings demonstrate that mitochondrial populations do not converge on a uniform bioenergetic endpoint during aging, but rather follow highly compartmentalized, tissue-specific spatiotemporal trajectories *in vivo*.

## 1 Introduction

Mitochondria act as the central hubs of cellular energy transduction, dynamically remodeling their structure and function to maintain metabolic homeostasis under age-related stress. This plastic response encompasses fission– and fusion-driven reorganization, fluctuations in mitochondrial membrane potential (ΔΨ*m*), and rigorous quality control through mitophagy and biogenesis [1]. During chronological aging, this quality control fails, leading to a profound decline in mitochondrial integrity. This age-associated dysfunction, recognized as a key hallmark of aging, disrupts cellular and tissue homeostasis through aberrant morphology, fragmented network dynamics, mitochondrial DNA mutations, elevated reactive oxygen species production, and bioenergetic collapse [1–3]. Ultimately, systemic mitochondrial decay acts as a critical upstream trigger for cellular senescence, chronic inflammation, and age-related tissue decline, culminating in organismal death [3].

During aging, the physical geometry and spatial topology of mitochondria undergo profound alterations. These changes often include a morphological shift from elongated, interconnected networks toward shorter, punctate spheres, driven by imbalanced fission-fusion kinetics, leading to network fragmentation and pathological swelling [4]. *Caenorhabditis elegans* (*C. elegans*) has been widely used to investigate these age-associated changes because its mitochondrial dynamics can be readily manipulated through genetic and environmental perturbations. These include genetic mutants with compromised fission-fusion machinery, such as fission-deficient *drp-1* (human *DRP1/DNM1L* ortholog), fusion-deficient *eat-3* (human *OPA1* ortholog), and fusion-deficient *fzo-1* (human *MFN1/MFN2* ortholog) mutants [5], as well as physiological stressors including heat stress [6], oxidative stress, and starvation/nutrient deprivation [7]. These targeted genetic and environmental perturbations can accelerate or mimic mitochondrial dysfunction, including loss of ΔΨ*m* and reduced bioenergetic efficiency, potentially leading to a metabolic shift from oxidative phosphorylation toward glycolysis [7].

A fundamental question is whether mitochondrial aging follows a coordinated organism-wide pattern or distinct tissue-specific trajectories. Growing evidence supports tissue-specific aging, where distinct tissues undergo divergent, adaptive metabolic reprogramming to meet unique physiological demands and microenvironmental cues [8]. Because mitochondria are central to energy production, metabolic regulation, and stress responses, they are likely key drivers and indicators of this tissue-specific aging. However, how mitochondrial states change across tissues during aging within the same living organism remains poorly understood.

A major barrier is the lack of methods that can resolve mitochondrial states across tissues *in vivo*. Organism– or organ-averaged readouts, including bulk tissue respirometry [9], quantitative PCR for mtDNA copy number estimation [10], and plate-reader-based ATP assays [11], fundamentally obscure tissue-, cell-type-, and sub-cellular compartment-specific heterogeneity. Evaluating ΔΨ*m* across distinct tissues remains one of the most accessible strategies for profiling localized metabolic activity. This is routinely accomplished by using classic, commercially available fluorophores, such as MitoTracker Red CMXRos [12], Rhodamine 123, JC-1, and tetramethylrhodamine derivatives such as Tetramethylrhodamine ethyl ester (TMRE) and tetramethylrhodamine methyl ester (TMRM) [13]. However, these conventional probes are prone to misinterpretation, as solely relying on uncalibrated fluorescent signal intensity introduces systematic bias during acquisition and analysis, eventually skewing measurements of mitochondrial activities [13]. While novel mitochondrial probes may mitigate these specific optical limitations [14–16], many of them are not widely accessible and have been validated mainly *in vitro* at the cellular level, leaving their efficacy and performance unproven at the organismal scale.

Fluorescence lifetime imaging microscopy (FLIM) provides a complementary readout by capturing time-resolved fluorescence decay signals generated by repeated excitation from a time-modulated laser source, rather than relying on fluorescence intensity alone [17]. While the emission spectrum of a given fluorophore may shift in response to environmental perturbations, such as alterations in solvent polarity, these spectral variations are typically subtle and remain undetectable when measuring integrated fluorescence intensity through a standard optical bandpass filter. Conversely, fluorescence lifetimes are highly sensitive to the local microenvironment and often exhibit pronounced variations driven by changes in pH, calcium levels, viscosity, polarity, molecular binding interactions, or conformational states [18]. FLIM intrinsically obtains traditional fluorescence intensity signals, giving an additional orthogonal dimension of contrast in optical imaging. Because FLIM acquires both intensity and lifetime information, it has been successfully leveraged to resolve and unmix multiple spectrally overlapping fluorophores through spectrally resolved FLIM (sFLIM) strategies [19], to map intracellular metabolic states via endogenous autofluorophores such as NADH and FAD [20], and to study cell metabolism in various organoid models for drug discovery and disease understanding [21]. To further extend these capabilities *in vivo*, two-photon fluorescence lifetime imaging microscopy (2p-FLIM) utilizes near-infrared (NIR) excitation rather than conventional visible-light single-photon laser sources. This approach offers enhanced optical penetration depth, intrinsic three-dimensional optical sectioning, minimized phototoxicity, and sub-micron spatial resolution comparable to state-of-the-art confocal systems, establishing 2p-FLIM as a well-suited tool for tissue-level biological research [22].

Here, we leverage 2p-FLIM to perform tissue-resolved, multi-parametric mitochondrial profiling at sub-cellular resolution within living, intact *C. elegans*. As a prominent, genetically tractable model organism, *C. elegans* has yielded foundational insights into the conserved regulatory mechanisms governing aging, neurobiology, genetics, and metabolism [23]. Its invariant cell lineage and stereotyped anatomy enable the same cells and tissues to be identified reproducibly across animals, providing a unique advantage for tissue– and cell-resolved analysis *in vivo* [23]. Utilizing the classic, mitochondria-targeted membrane potential probe TMRM, we demonstrate that orthogonal structural and functional datasets can be simultaneously extracted from 2p-FLIM measurements of the living animals across their lifespan. Specifically, while calibrated two-photon excitation fluorescence (TPEF) intensity tracks alterations in mitochondrial energetics, the corresponding fluorescence lifetime captures subtle, tissue-dependent variations in the intraorganellar microenvironment. By coupling this imaging modality with a novel segmentation framework termed Mitochondria Segmentation with Locally Iterative Thresholding (MitoSLIT), machine learning, and quantitative morphometric analysis, we map mitochondrial network architectures and structural reorganization during aging and under various mutant and stress conditions. Using this approach, we reveal: (1) a remarkably diverse, tissue-specific landscape of mitochondrial states and aging trajectories across somatic tissues; (2) to the best of our knowledge, the first *in vivo* validation and characterization of neuronal mitochondrial staining within living, intact nematodes and the compartment-specific changes in mitochondrial states, morphology, and redox balance within individual neurons during aging; and (3) distinct relationships between physiological aging and mitochondrial remodeling induced by stress or altered fission-fusion dynamics. Finally, we identify sample-mounting-induced hypoxia during imaging as a critical confounding factor that substantially alters mitochondrial morphology and bioenergetic readouts, highlighting a systematic artifact that must be rigorously controlled or avoided during *in vivo* mitochondrial assays.

## 2 Results

### 2.1 Optimized *in vivo* 2p-FLIM reveals mitochondrial heterogeneity across tissues in young adult *C. elegans*

Mitochondrial membrane potential (ΔΨ*m*) has been routinely and widely used as a functional indicator of the local mitochondrial bioenergetic state. Among several commercially available fluorescent membrane-potential dyes such as Rhodamine 123, TMRE, and TMRM, TMRM exhibits comparatively low nonspecific binding and respiratory inhibition [24]. Nevertheless, like other membrane-potential-sensitive dyes, TMRM is susceptible to experimental artifacts and misinterpretation if the staining conditions, including dye concentration and incubation time, are not rigorously verified [13]. To optimize the experimental parameters, we tested various TMRM staining conditions under 2p-FLIM microscopy by following a standardized framework [13], evaluating concentrations from tens to hundreds of nM across incubation windows ranging from 30 minutes to overnight. We found that 1.5-hour staining on OP50 plates containing 200 nM TMRM provided a sufficiently high signal-to-noise ratio. In transgenic wild-type adult worms expressing mito::GFP in their body-wall muscles (*zcIs14 [myo-3::GFP(mit)]*), the TMRM signal was strongly colocalized with GFP-labeled mitochondria and showed a strong positive linear relationship across different age groups (Pearson’s correlation coefficient = 0.956 ± 0.06, mean ± SD; Fig. 1A), supporting consistent mitochondrial localization of TMRM under the staining condition in living *C. elegans*. We validated the ΔΨ*m* sensitivity and specificity of the TMRM staining condition using FCCP, a potent mitochondrial protonophore uncoupler. The pharmacological dissipation of ΔΨ*m* by FCCP markedly reduced calibrated TMRM intensity (see “Calibrated TMRM intensity” in Methods for details) (Fig. S1), confirming the sensitivity of the staining regimen to mitochondrial depolarization.

**Fig. 1:**
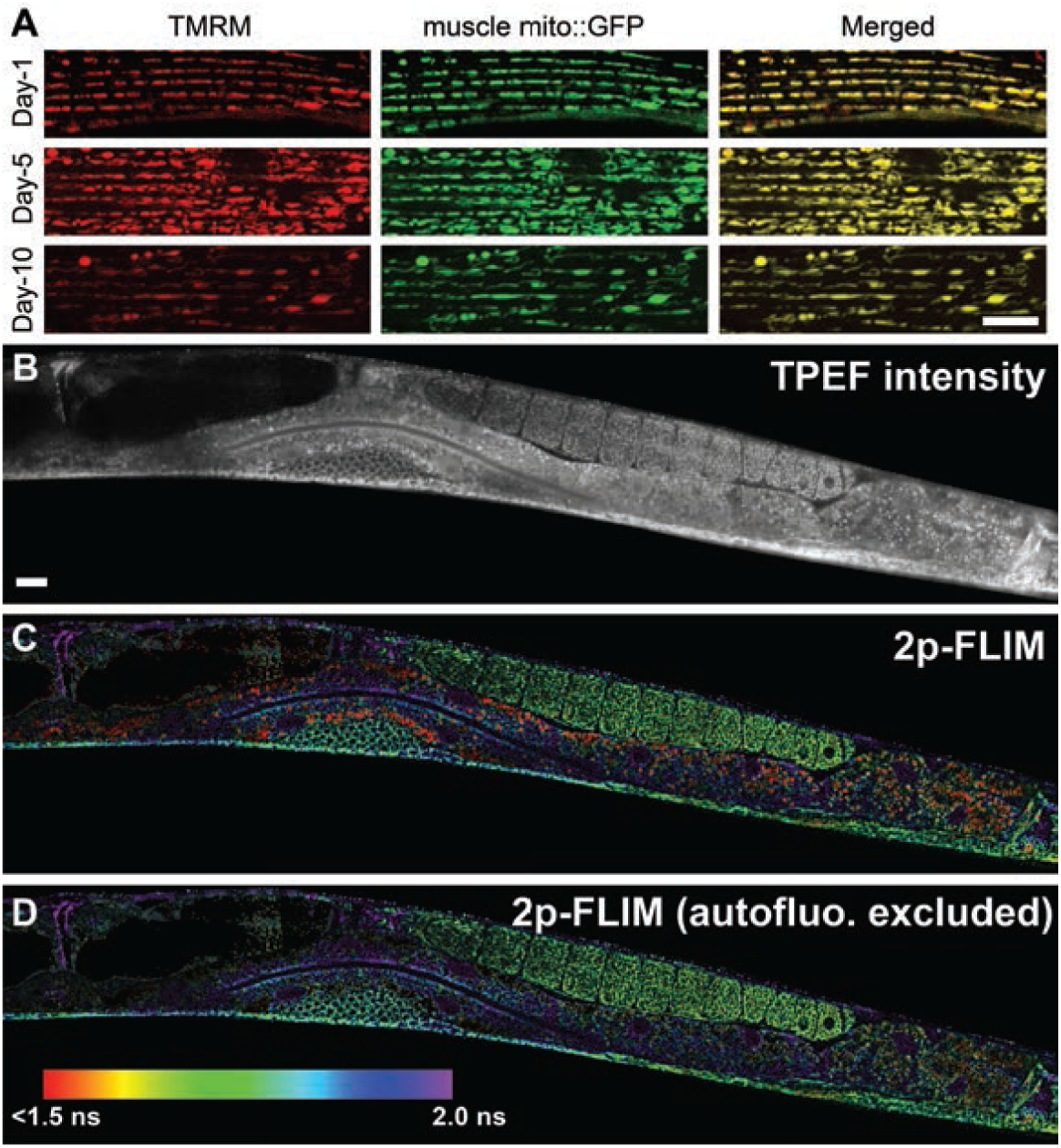
2p-FLIM *in vivo* imaging of *C. elegans* with TMRM staining. (A) Representative images showing colocalization of muscle mito::GFP (green) and TMRM (red) in day-1, day-5 and day-10 *C. elegans* (*zcIs14 [myo-3::GFP(mit)]*); merged images are shown in yellow. Images of (B) TPEF intensity, (C) 2p-FLIM, and (D) 2p-FLIM after excluding autofluorescence (<1.3 ns) from a day-1 adult wild-type worm stained with TMRM. The color scale indicates lifetimes from <1.5 ns to 2.0 ns. Scale bars, 10 *µ*m.

Having optimized and validated the staining conditions, we next applied TMRM-based 2p-FLIM to multiple organs and tissues (see Methods for details), enabling the high-resolution detection of mitochondrial networks in live animals (Fig. 1B-D). Beyond variations in TMRM fluorescence intensity, distinct fluorescence lifetime (*τ*) signatures across diverse tissues reveal significant heterogeneity in the mitochondrial microenvironment (Fig. 1C). In the intestine, prominent autofluorescent gut granules, often described as lipofuscin-like or age-pigment-associated autofluorescence [25], were detected within the TMRM emission channel. However, they exhibit a characteristic short fluorescence lifetime (≤1.3 ns) [26] and thus can be effectively excluded through lifetime gating (Fig. 1D). Collectively, these results demonstrate that the combination of 2p-FLIM and the commercially available dye TMRM can reliably detect mitochondrial localization, morphology, and tissue-specific bioenergetic heterogeneity in live *C. elegans*. Our data further reveal that even in chronologically “young and healthy” 1-day adult animals, mitochondria manifest tissue-specific energetic states, reflecting a high degree of functional specialization prior to age-related decline.

### 2.2 Short-term hypoxia is sufficient to perturb mitochondrial metabolic states and morphological patterns *in vivo*

Mitochondrial bioenergetic and morphological states are highly sensitive to oxygen availability, with hypoxia rapidly perturbing respiratory function and redox balance. This sensitivity is particularly relevant to *in vivo* imaging, as conventional *C. elegans* immobilization between an agarose pad and glass coverslip can restrict gas exchange and induce hypoxic stress, potentially producing mitochondrial changes that may be misinterpreted as physiological or experimental effects. We observed that after 30-40 minutes under a glass coverslip, mitochondrial networks underwent pronounced morphological remodeling, transitioning from tubular, filamentous networks to irregularly dispersed punctate structures. In contrast, replacing the glass coverslip with a gas-permeable coverslip preserved an elongated and densely organized mitochondrial architecture over a 90-minute imaging window (Fig. 2A,B).

**Fig. 2:**
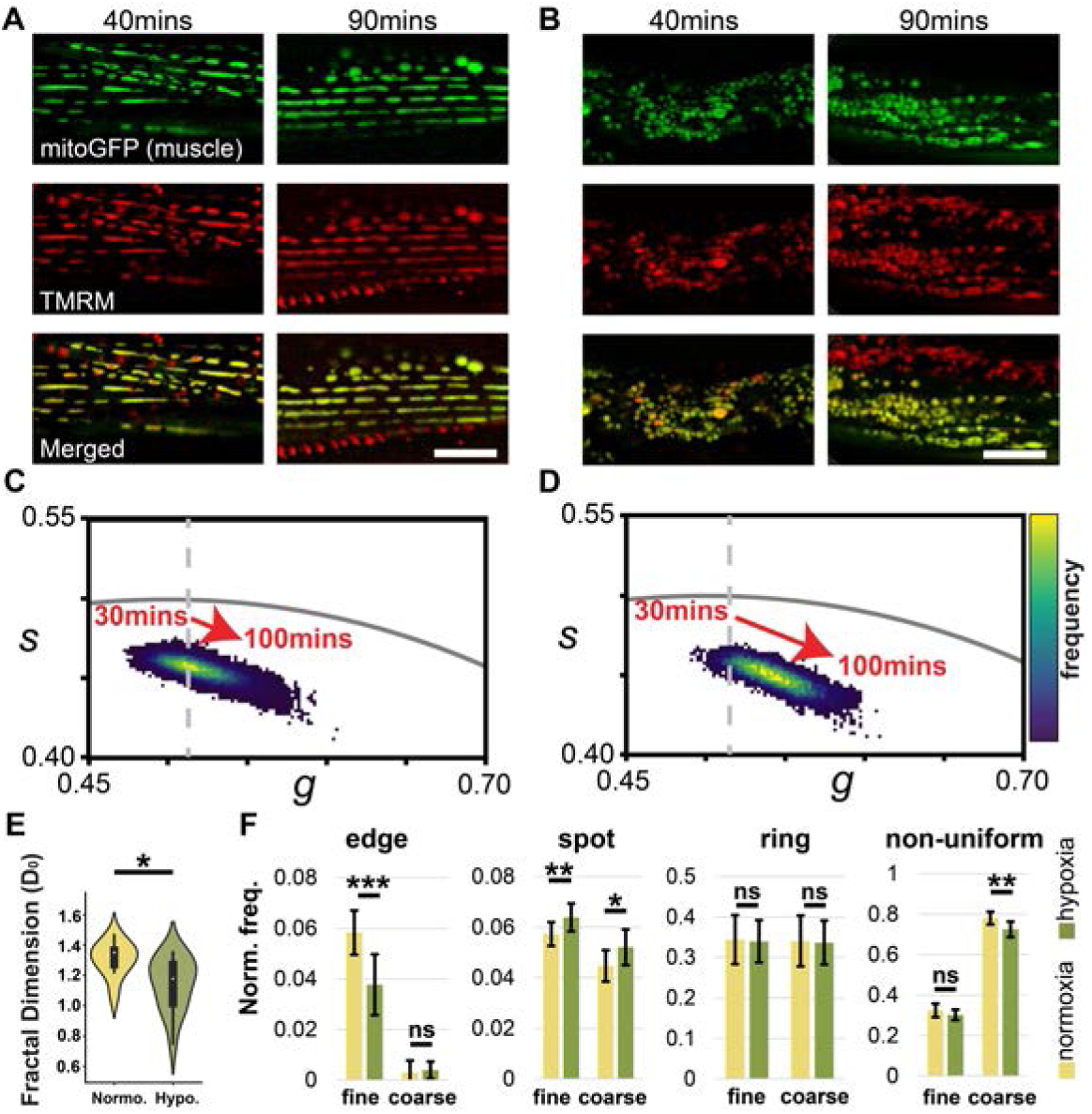
Hypoxia perturbs mitochondrial metabolic states and morphological patterns. (A, B) Representative body-wall muscle mitochondrial images from day-1 *Pmyo-3::GFP(mit)* worms showing mito::GFP (top), TMRM (middle), and merged overlays (bottom) after 40 min and 90 min of mounting under (A) a gas-permeable polymer coverslip (normoxia) or (B) a sealed glass coverslip (hypoxia). (C, D) Phasor plots of TMRM signal from day-1 *Pmyo-3::GFP(mit)* worms acquired over a 100-min imaging window under (C) normoxia or (D) hypoxia conditions. (E, F) Morphological pattern changes under hypoxia. Quantification of body-wall muscle mitochondrial organization under normoxia versus hypoxia using (E) fractal analysis. (F) Rotation-invariant local binary pattern (LBP) features, reported as normalized mean frequencies for edge, spot, ring, and non-uniform patterns at fine and coarse spatial scales. Error bars represent standard deviation. Significance was assessed using a Welch’s t-test; ns, not significant; *p* < 0.05 (*), *p* < 0.01 (**), and *p* < 0.001 (***). Scale bars, 10 *µ*m.

To quantitatively evaluate mitochondrial morphological transitions under these distinct environments, we applied higher-order morphometric and textural pattern analysis using box-counting fractal dimension and rotationally invariant Local Binary Patterns (LBP), which capture global space-filling complexity and scale-dependent local texture organization, respectively (details are provided in the subsections of “Mitochondrial morphological analysis” in Methods). Hypoxia significantly reduced the mitochondrial capacity dimension (*D*0), consistent with diminished space-filling complexity (Fig. 2E). LBP analysis further revealed a loss of fine-scale edge-like features and an increase in spot-like patterns, supporting a transition toward a more punctate organization (Fig. 2F).

Beyond this morphological remodeling, hypoxia altered the mitochondrial fluorescence lifetime signatures. Under glass coverslips, the phasor cluster centroid followed a marked and continuous trajectory over the 100-minute acquisition period, consistent with a progressive shifting and reduction in TMRM fluorescence lifetime (Fig. 2D). In contrast, gas-permeable coverslips produced a stable phasor distribution throughout imaging (Fig. 2C), indicating that oxygen limitation perturbs the local mitochondrial microenvironment.

Collectively, these data demonstrate that acute hypoxic stress alters both mitochondrial spatial organization and TMRM fluorescence lifetime signatures. Mitochondria changed from dense, tubular networks to irregular, low-density void patterns even in young wild-type animals, demonstrating that these changes can arise from hypoxia induced by imaging conditions rather than from confounding biological factors such as aging or genetic mutations. Because these hypoxia-driven systemic artifacts can profoundly distort both TMRM signal interpretations and mitochondrial networking metrics, they must be rigorously controlled and minimized in all *in vivo* mitochondrial assays. Consequently, all subsequent experiments were performed using gas-permeable coverslips to minimize hypoxia-induced perturbations.

### 2.3 Aging reshapes mitochondrial states in a tissue-dependent manner

Having established tissue-resolved mitochondrial differences in young adults and minimized sample-mounting-induced hypoxia, we next asked whether these mitochondrial states follow common or distinct trajectories during aging. To address this question, we examined mitochondrial 2p-FLIM signals in the hypodermis, gonad, and pharynx of day-1, day-5, and day-10 wild-type worms (Fig. 3). From the same 2p-FLIM datasets, we first analyzed calibrated TMRM fluorescence intensity as the conventional ΔΨ*_m_*-associated readout. Across all three tissues, calibrated TMRM fluorescence intensity decreased monotonically with age (Fig. S3A), indicating reduced mitochondrial TMRM retention and age-associated decline in ΔΨ*_m_*. This is consistent with the conventional TMRM-based paradigm, in which reduced TMRM intensity reflects mitochondrial depolarization under non-quenching mode [13].

**Fig. 3:**
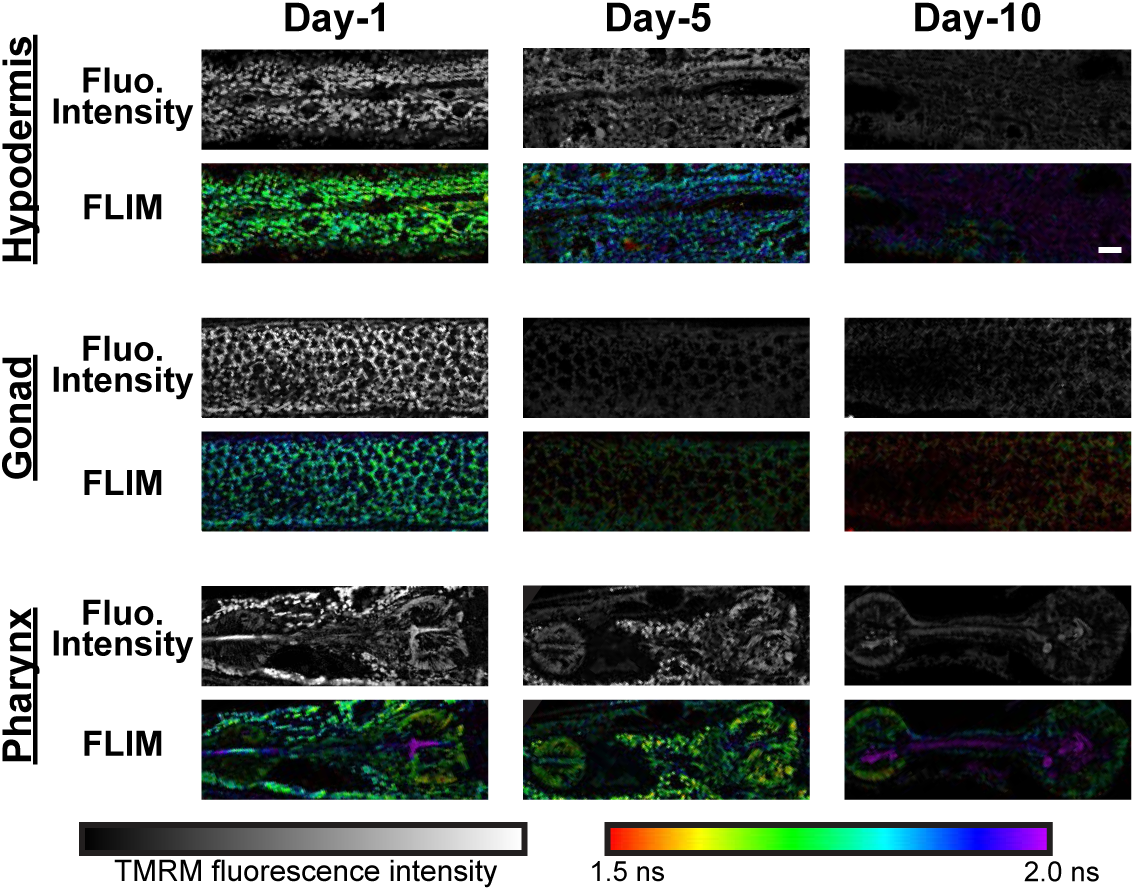
Tissue-dependent fluorescence intensity and lifetime changes during aging. Representative two-photon TMRM fluorescence intensity and FLIM images from the hypodermis, gonad, and pharynx of day-1, day-5, and day-10 adult *C. elegans*. Grayscale images indicate TMRM fluorescence intensity, and pseudocolor images indicate fluorescence lifetime. FLIM color scale, 1.5-2.0 ns. Scale bars, 10 *µ*m.

Although TMRM fluorescence intensity decreased with age across all three tissues, the kinetics of this decline were tissue-specific (Fig. 3 and Fig. S3A). In the hypodermis, intensity diminished progressively, where the signal was halved by day 5 and dropped to ∼11% by day 10. In contrast, the gonad exhibited an early, precipitous loss of signal. By day 5, fluorescence intensity dropped by nearly an order of magnitude, rendering the characteristic honeycomb-like pattern nearly indistinguishable. This sharp reduction coincides with the onset of reproductive senescence and likely reflects the large-scale germline remodeling associated with the exhaustion of self-fertility [27]. By day 10, the gonadal signal reached near-baseline levels (∼5% of day 1), while the remaining mitochondrial structures exhibited a more porous and fragmented reticular architecture. In the pharynx, the signal was restricted to the muscle of the terminal bulb after excluding bright confounding signals from residual *E. coli* within the central lumen. The intensity also declined with age, reaching approximately 30% of the day-1 level by day 5 and 16% by day 10, indicating a substantial early reduction followed by a more moderate additional decrease. This age-dependent loss of pharyngeal mitochondrial signal occurs in the context of previously reported functional and structural decline of the aging *C. elegans* pharynx, including sarcopenia-like changes [28].

We next asked whether fluorescence lifetime could reveal tissue-specific changes that were not captured by conventional intensity measurements. Strikingly, in contrast to the monotonic decline in intensity, FLIM signatures exhibited tissue-specific trajectories and heterogeneity during aging (Fig. 3 andFig. S3B). We compared pooled pixel-level lifetime distributions and per-image mean lifetimes to assess distributional remodeling together with image-level shifts. In the hypodermis, fluorescence lifetimes increased progressively, as demonstrated by a rightward shift in the pooled distribution and substantial increases in per-image mean lifetime at day 5 and day 10. This shift was accompanied by distribution broadening and the emergence of a high-lifetime tail, indicating increased lifetime heterogeneity. Conversely, the gonad showed age-dependent lifetime shortening, with a leftward shift in the pooled distribution, a significant decrease in mean lifetime, enrichment of low-lifetime components, and loss of higher-lifetime populations. The pharynx showed comparatively modest changes. Although its pooled distribution and mean lifetime shifted slightly upward by day 10, the distributions remained highly overlapping across ages.

Together, these results indicate that while the age-dependent decline in TMRM intensity is consistent with reduced ΔΨ*m*, FLIM lifetime captures an additional, tissue-specific dimension of mitochondrial aging, with lifetime trajectories and distributional heterogeneity varying markedly across tissues. This raised the question of how intensity– and lifetime-associated features are jointly reorganized within mitochondrial populations across tissues and ages, prompting an integrated pixel-level analysis that preserves both phasor-intensity information and spatial context.

### 2.4 MitoSLIT reveals coordinated bioenergetic and structural remodeling of mitochondria during aging

A prerequisite for this integrated analysis was reliable identification of mitochondrial pixels across tissues and ages. The pronounced age-dependent decline in TMRM intensity, together with spatial variations in local background and mitochondrial signal contrast, presented a challenge for conventional segmentation. Standard intensity-based approaches, such as Otsu’s global thresholding [29], can preferentially exclude dim mitochondrial structures, making a single global threshold unsuitable for comparisons across tissues and ages. To overcome this limitation, we developed a new segmentation method, Mitochondria Segmentation with Locally Iterative Thresholding (MitoSLIT), which identifies mitochondrial structures based on local intensity contrast rather than absolute fluorescence intensity (details are provided in the section “Mitochondria Segmentation with Locally Iterative Thresholding” in Methods and Fig. S4). Using a fixed parameter set, this local, probability-based strategy detected both bright and dim mitochondrial structures across tissues and ages despite differences in signal intensity and background levels (Fig. S5).

We next converted the time-domain lifetime decay information into phasor components (S and G) [30] and integrated FLIM phasor components and calibrated TMRM intensity across ∼3×10^7^ MitoSLIT-identified mitochondrial pixels. Following fluorescein-based calibration and laser-power normalization, each pixel was represented by a three-dimensional phasor-intensity feature vector (*G, S, I*calibr). Mitochondrial pixels from the hypodermis, gonad, and pharynx across aging and perturbation conditions were pooled and subjected to unsupervised k-means clustering after feature standardization (Z-score). Cluster-validation analyses supported eight phasor-intensity mitochondrial states (Fig. 4A and Fig. S6), while UMAP [31] visualization showed that these clusters occupied largely distinct regions of the feature space. Each mitochondrial pixel was subsequently assigned to one of the eight states and mapped back to its original spatial location. We then examined these clustering-defined states in day-1, day-5, and day-10 wild-type animals to characterize tissue-specific aging trajectories. We found that, although local brightness variation was present, intensity alone primarily reported changes in signal magnitude and provided a comparatively coarse view of mitochondrial remodeling (Fig. 4B). In contrast, the corresponding phasor-intensity cluster maps provided a spatially resolved visualization of mitochondrial state heterogeneity that was not apparent from intensity alone (Fig. 4C).

**Fig. 4:**
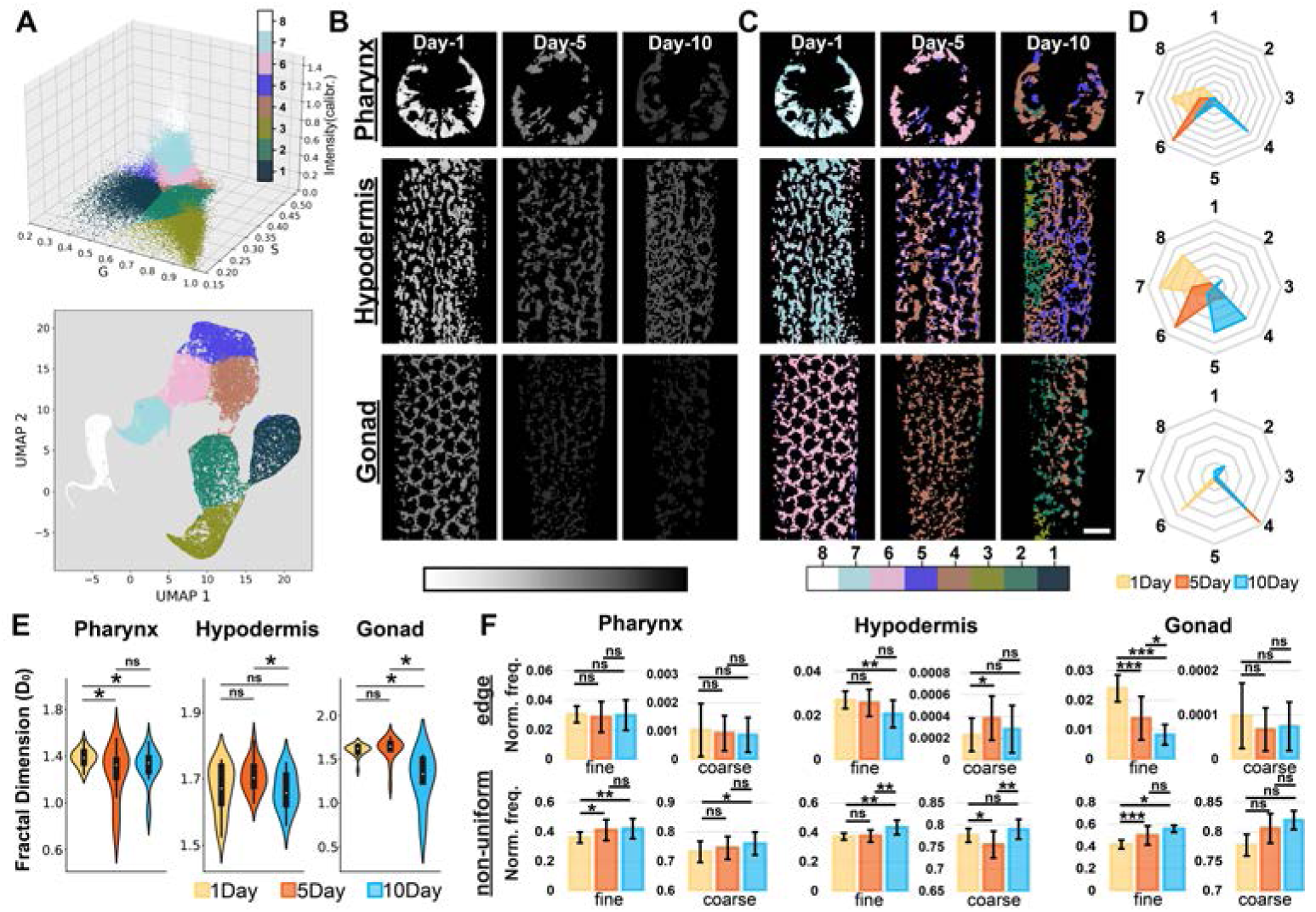
Coordinated bioenergetic and structural remodeling of mitochondria during aging. (A) Mitochondrial pixels were clustered into eight states based on integrated fluorescence lifetime phasor coordinates (*G, S*) and calibrated fluorescence intensity (I*_calibr_*). The upper plot shows the distribution of clustered mitochondrial pixels in G-S-I*_calibr_* space, and the lower plot shows the corresponding UMAP projection of the clustering pixel populations. (B) Representative TMRM fluorescence intensity images from the pharynx, hypodermis, and gonad of day-1, day-5, and day-10 adult *C. elegans*. (C) Corresponding clustering images showing spatial distributions of the eight mitochondrial states across tissues and ages. Scale bars, 10 *µ*m. (D) Radar plots summarizing age-dependent changes in mitochondrial state composition for each tissue. (E) Fractal dimension analysis of mitochondrial patterns across aging. Violin plots show image-level fractal dimension (*D*0). (F) Local binary pattern analysis of mitochondrial texture features at fine and coarse spatial scales. Bar plots summarize normalized frequencies of edge-like and non-uniform texture features. Clustering in (A) was performed on ∼3×10^7^ mitochondrial pixels from 269 images spanning three tissues and multiple experimental conditions. For WT aging analyses, sample sizes at day 1, day 5, and day 10 were *n* = 15, 16, and 11 images for hypodermis, *n* = 19, 16, and 6 for gonad, and *n* = 24, 16, and 21 for pharynx, respectively. Bars indicate mean ± SD. Pairwise comparisons were performed using two-tailed Welch’s t-test. ns, not significant; *p* < 0.05 (*), *p* < 0.01 (**), and *p* < 0.001 (***).

In the hypodermis, day-5 animals showed a more diffuse redistribution of mitochondrial states throughout the tissue, whereas day-10 animals exhibited more pronounced subregional enrichment of distinct states, suggesting that age-related mitochondrial changes became concentrated in specific regions of the hypodermis by day 10. In the pharynx, the internal regions of the terminal bulb exhibited distinct patterns of phasor-intensity cluster assignment from those in the surrounding muscle, revealing localized heterogeneity that was obscured by conventional intensity-based imaging and suggesting that age-related mitochondrial remodeling is spatially organized within the tissue. In contrast, gonadal state redistribution appeared comparatively uniform across the tissue, consistent with a more coordinated tissue-wide shift in state composition (Fig. 4C,D). Taken together, these results indicate that mitochondrial aging does not simply follow a uniform decline in ΔΨ*m*, but instead involves spatially organized and tissue-specific redistribution of mitochondrial states.

We next asked whether this mitochondrial state remodeling was accompanied by changes in mitochondrial architecture. We therefore analyzed MitoSLIT-derived mitochondrial patterns using fractal dimension and LBP-based texture features. At the global architectural level, fractal dimension declined with age across all three tissues, although the magnitude and timing differed among tissues, indicating a loss of highly reticulated, space-filling mitochondrial organization by day 10 (Fig. 4E). The results of LBP-based analysis show a further layer of highly tissue-specific morphological restructuring (Fig. 4F and Fig. S7). In the hypodermis, reduced fine-scale edge-like features together with increased non-uniform texture indicated a shift from an evenly reticulated local organization toward a more heterogeneous mitochondrial pattern (Fig. 4F). The gonad underwent the most pronounced textural remodeling, with reduced edge-like features and increased spot-like and non-uniform patterns, consistent with a transition from continuous tubular structures toward more fragmented and punctate organization (Fig. 4F and Fig. S7). The concurrent loss of coarse-scale ring-like features further reflected deterioration of the characteristic honeycomb-like arrangement in young gonads (Fig. S7). In contrast, pharyngeal edge-like features remained comparatively stable, while changes at broader spatial scales indicated the development of a more irregular, patchy organization (Fig. 4F and Fig. S7).

Together, these analyses suggest that mitochondrial aging involves concurrent remodeling of mitochondrial state and structural organization.

### 2.5 Multiparametric mapping reveals tissue-specific, aging-associated mitochondrial remodeling trajectories

To determine whether this integrated aging signature was specific to physiological aging or could be phenotypically recapitulated by defined stress conditions and mitochondrial defects, we applied our multiparametric framework to animals exposed to exogenous stressors or carrying genetic perturbations that accelerate or mimic distinct facets of mitochondrial aging. Specifically, we profiled wild-type animals subjected to thermal stress, oxidative stress, and starvation, alongside mutant strains deficient in mitochondrial fission-fusion machinery (*drp-1*, *eat-3*, and *fzo-1*) (Fig. 5A). By integrating mitochondrial state profiles with morphology-derived structural metrics, we computed pairwise similarity matrices across aging, mutant, and stress cohorts for each tissue (Fig. 5B).

**Fig. 5:**
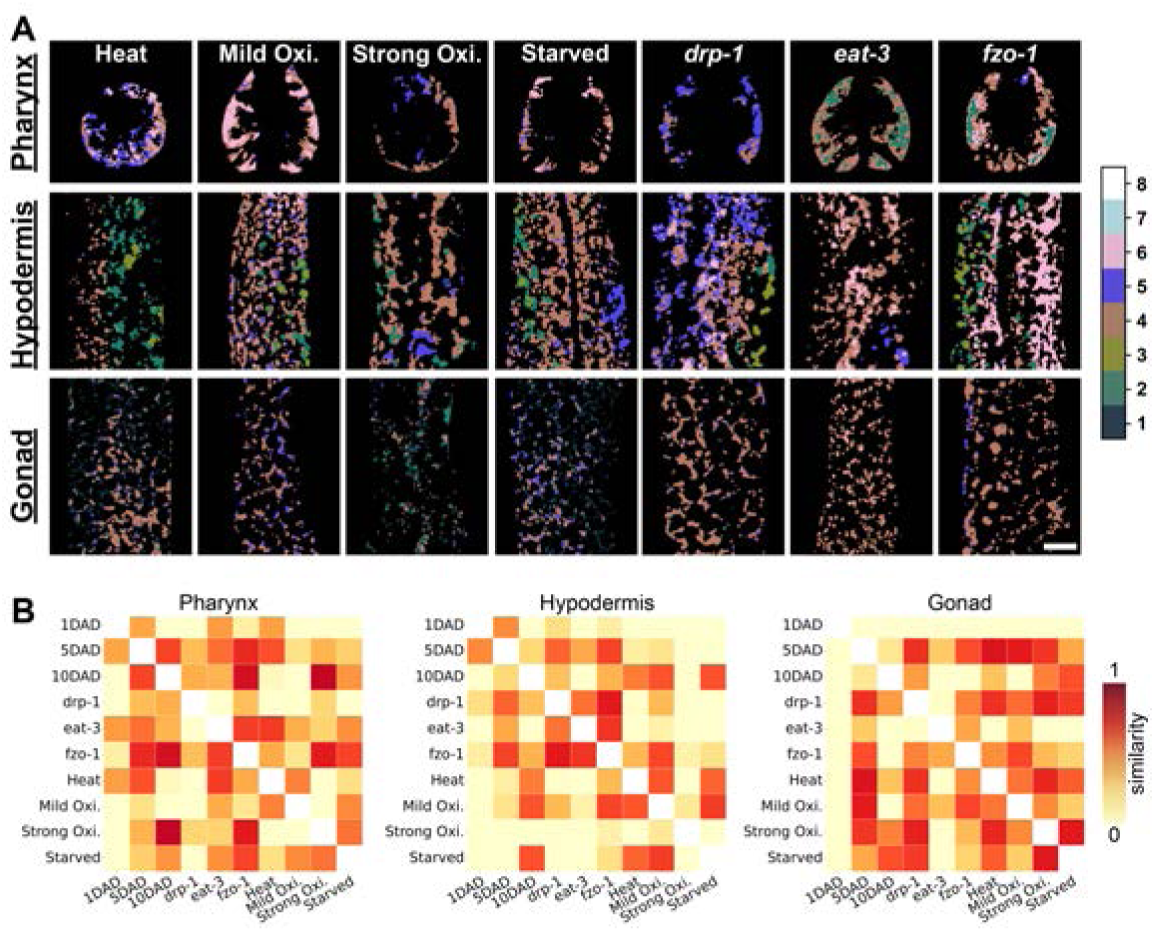
Similarity analysis compares aging-associated mitochondrial remodeling with stress and mitochondrial dynamics perturbations. (A) Clustered mitochondrial images from the pharynx, hypodermis, and gonad under acute stress conditions and mitochondrial dynamics perturbations, including heat stress, mild oxidative stress, strong oxidative stress, starvation, and genetic perturbation of mitochondrial fission/fusion regulators *drp-1*, *eat-3*, and *fzo-1*. Scale bars, 10 *µ*m. (B) Conditioned similarity matrices comparing mitochondrial phenotypes across aging, stress, and genetic perturbation conditions within each tissue. Similarity was calculated using integrated mitochondrial state composition and morphology-derived features. Warmer colors indicate greater similarity between conditions. For the aging groups (1-, 5-, and 10-day-old adults, respectively), *n* = 21, 16, and 11 animals for the hypodermis; *n* = 22, 16, and 6 for the gonad; and *n* = 28, 16, and 21 for the pharynx. For the perturbation groups (*drp-1*, *eat-3*, *fzo-1*, heat stress, mild oxidative stress, strong oxidative stress, and starvation, respectively), *n* = 14, 8, 17, 16, 8, 8, and 16 animals for the hypodermis; *n* = 14, 8, 16, 13, 8, 9, and 16 for the gonad; and *n* = 14, 8, 16, 16, 8, 9, and 16 for the pharynx. Each *n* represents one independently imaged animal.

The resulting tissue-specific similarity matrices revealed that exogenous stressors and genetic mutations did not uniformly recapitulate a singular aged mitochondrial state. Across all examined tissues, most perturbations exhibited low similarity to the day-1 baseline, indicating substantial deviation from a youthful mitochondrial state. However, the specific chronological aging stage mirrored by each perturbation was highly dependent on both tissue type and perturbation class (Fig. 5B). The most pronounced convergence occurred in the gonad, where nearly all stress and mutant conditions shifted away from the day-1 profile and aligned closely with the day-5 state. This transition coincides with the onset of reproductive aging. By day 5, hermaphrodites exhibit a sharp decline in reproductive output, falling to approximately 6% of their peak progeny production rate as they approach the end of their self-fertile period [27]. By day 10, following the cessation of reproduction, the gonadal mitochondrial landscape more closely resembled phenotypes induced by severe oxidative stress and starvation. These findings are consistent with established links between mitochondrial dynamics and metabolism, oocyte quality, and reproductive aging in *C. elegans* [32, 33], as well as extensive gonadal remodeling during programmed reproductive death, a self-destructive quasi-program that accelerates systemic organismal aging but can be suppressed via germline ablation [34].

In the hypodermis, the alignment between acute perturbations and physiological aging was more condition-specific. Phenotypes driven by disrupted mitochondrial dynamics (e.g., *drp-1* and *fzo-1* mutants) closely resembled the mid-life day-5 profile rather than the late-life day-10 profile. This finding suggests that an intermediate stage of hypodermal aging shares phenotypic features with altered mitochondrial fission-fusion dynamics. Conversely, thermal stress, mild oxidative stress, and starvation aligned primarily with late-aging signatures. This day-10 hypodermal profile is consistent with reported age-dependent fragmentation of mitochondrial networks, respiratory decline, and attenuation of the heat-shock response in *C. elegans* [6]. In contrast to the more stage-associated shifts in the hypodermis and the broad convergence observed in the gonad, the pharynx displayed a more progressive, overlapping similarity trajectory. At both day 5 and day 10, pharyngeal profiles shared features with both fission-fusion defects (such as *fzo-1*) and oxidative stress conditions, suggesting that age-associated mitochondrial remodeling in the pharynx follows a more gradual and overlapping trajectory rather than discrete state transitions. Together, these comparisons show that defined mitochondrial perturbations recapitulate selected features of physiological aging in a tissue-dependent manner, but no single stress or genetic defect reproduces a universal aged mitochondrial state across tissues.

### 2.6 Age-associated mitochondrial remodeling in single neurons

Remarkably, our imaging protocol successfully resolved mitochondria within single neurons by employing the same TMRM concentration with a slightly longer incubation window (∼3 hours). To verify that the observed TMRM-positive structures localized within neuronal somata, we examined transgenic animals expressing cytosolic GFP in a defined subset of neurons, including the RIC and RIM interneurons [35]. Sum-intensity z-projections isolated from z-slices containing RIC/RIM GFP signals revealed clearly discernible TMRM-labeled mitochondria (Fig. 6A). To independently confirm whether the TMRM-positive structures within neurons were mitochondria, we examined a transgenic strain expressing mitochondria-outer-membrane targeted GFP (mito::GFP) specifically within the AIY interneurons [36]. Sum-intensity z-projections demonstrated strong spatial overlap between TMRM fluorescence and the localized mito::GFP signal, confirming that mitochondrial networks within individual neurons can be unambiguously resolved *in vivo* using our 2p-FLIM pipeline (Fig. 6B). Although this approach requires an additional neuronal marker, to the best of our knowledge, this constitutes the first demonstration of *in vivo* TMRM staining and mitochondrial visualization within individual neurons of *C. elegans*.

**Fig. 6:**
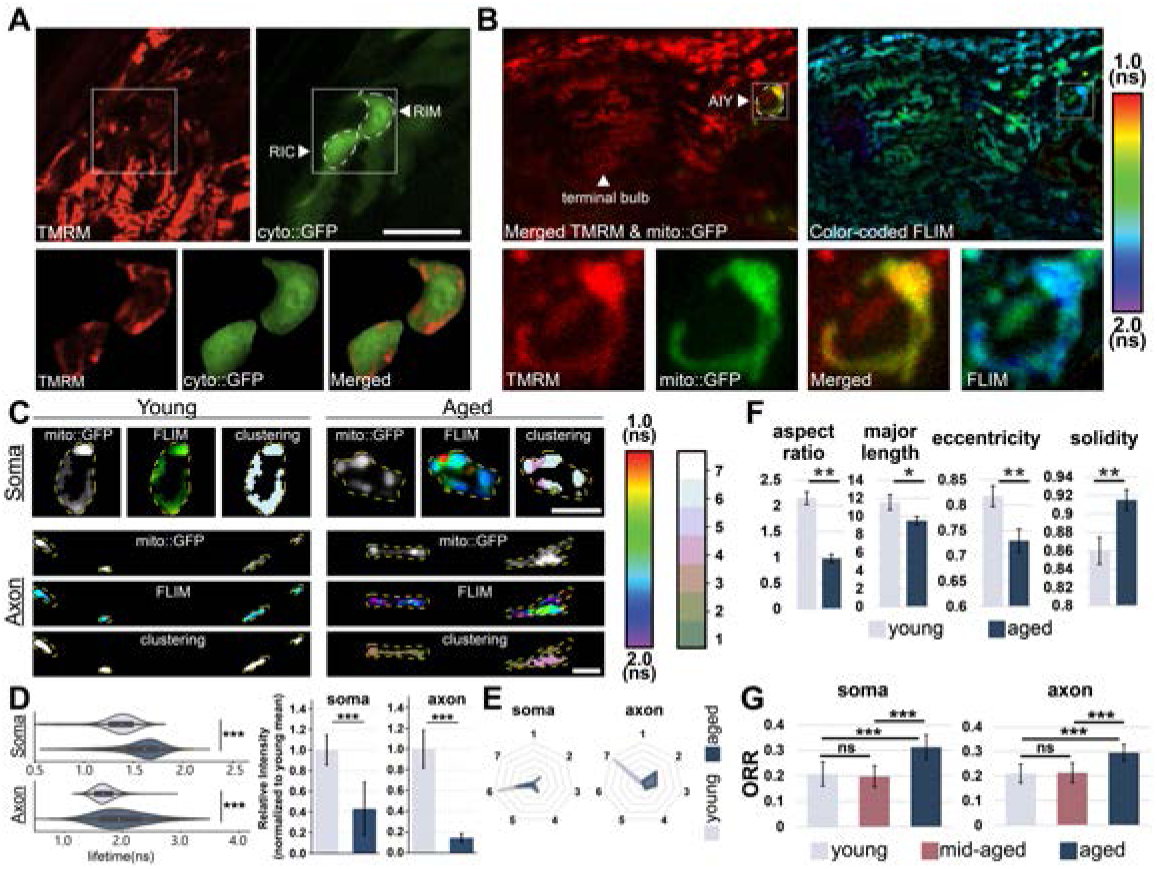
Single-neuron-specific TMRM-FLIM imaging reveals age-associated mitochondrial metabolic and morphological remodeling. (A) Representative images showing TMRM fluorescence in GFP-labeled RIC and RIM neurons. Cytosolic GFP was used to identify neuronal soma regions, and TMRM-positive structures were observed within the GFP-defined neuronal regions. Magnified views show TMRM, cyto::GFP, and merged signals within the selected region. (B) TMRM labeling in AIY neurons using the AIY-specific mito::GFP strain. TMRM fluorescence overlapped with mito::GFP, confirming mitochondrial labeling within genetically defined AIY neurons. Color-coded FLIM images show the TMRM fluorescence lifetime distribution within the AIY region, with magnified views of TMRM, mito::GFP, merged fluorescence, and FLIM signals. (C) Representative young and aged AIY neuronal mitochondria in soma and axon regions. Mito::GFP was used to delineate mitochondrial regions, and corresponding FLIM and clustered-mitochondria state images were generated within the GFP-defined mitochondrial masks. Yellow dashed outlines indicate segmented mitochondrial regions. (D) Quantification of mitochondrial TMRM fluorescence lifetime (left) and calibrated fluorescence intensity (right) in the AIY soma and axon of young and aged animals. Violin plots show the distributions of TMRM fluorescence lifetime, and bar graphs show calibrated TMRM intensity normalized to the corresponding day-1 (young) mean. Bars represent mean ± SD; n = 10 animals per group. (E) Radar plots summarizing age-dependent changes in clustered mitochondrial state composition in soma and axon regions. (F) Morphological analysis of neuronal mitochondria in the soma region showing age-associated changes in aspect ratio, major axis length, eccentricity, and solidity. Bars indicate mean ± SD. (G) Optical redox ratio (ORR) calculated as ORR = FAD / [FAD + NAD(P)H], in neuronal soma and axon regions across young (day-1), mid-aged (day-5), and aged animals (day-10). Bars indicate mean ± SD; n = 10 for the young and mid-aged groups and n = 11 for the aged group. Statistical comparisons were performed using two-tailed Welch’s t-test. ns, not significant; *p* < 0.05 (*), *p* < 0.01 (**), and *p* < 0.001 (***).

We next investigated age-associated mitochondrial remodeling within individual sub-neuronal compartments. To capture the full spatial topography of mitochondria distributed across the 3D neuronal architecture, we acquired z-stacks of TMRM-stained AIY neurons. Quantification of calibrated TMRM fluorescence intensity revealed a marked age-associated decline in both compartments. Relative to the corresponding day-1 mean, the normalized TMRM intensity decreased to ∼43% in the soma and ∼14% in the axon of aged animals (Fig. 6D right). Representative FLIM maps revealed an age-dependent shift toward longer TMRM fluorescence lifetimes across both cellular compartments (Fig. 6C), and quantitative profiling confirmed significantly longer lifetimes in aged animals than in their young counterparts (Fig. 6D left). The lifetime distribution violin plots also reveal compartment-specific remodeling (Fig. 6D left). While the soma exhibited a relatively simple rightward shift with age, the axonal compartment displayed pronounced distribution broadening, characterized by an enriched subpopulation with substantially longer lifetimes.

Unsupervised clustering analysis further supported this compartment-specific divergence. In young adult animals, neuronal mitochondrial populations in both the soma and axon clustered tightly within a relatively narrow phenotypic state. Conversely, aged neurons exhibited a marked population shift paired with expanded phenotypic heterogeneity (Fig. 6C,E, and Fig. S9). These transitions indicate that physiological aging drives an expansion of mitochondrial state heterogeneity within individual neurons. This phenotypic divergence was particularly pronounced in the axonal compartment, aligning with the heightened vulnerability of distal axonal mitochondria to age-related maintenance declines [37, 38].

The mitochondrial state shifts observed by TMRM imaging prompted us to assess whether neuronal mitochondria concurrently undergo structural remodeling. Unlike the dense, highly convoluted mitochondrial networks found in larger somatic tissues, AIY neuronal mitochondria are sparsely distributed and spatially discrete under high-magnification imaging, allowing transition from network-level texture analysis to a single-organelle morphometric framework. Using watershed segmentation [39] applied to MitoSLIT-defined regions in the images, individual mitochondrial boundaries were isolated and quantified across geometric descriptors. In the AIY neuronal soma, aging mitochondria exhibited significant reductions in aspect ratio, major axis length, and eccentricity, accompanied by increased solidity (Fig. 6F), indicating remodeling from elongated mitochondrial profiles toward shorter, rounder, and more compact structures.

To complement the mitochondrial state and structural remodeling observed during neuronal aging, we evaluated the optical redox ratio (ORR) in the AIY somatic and axonal regions of day-1, day-5, and day-10 animals. The compartment-specific ORR was quantified as the sum-intensity ratio of flavin adenine dinucleotide to total metabolic cofactors, expressed as [FAD / (NAD(P)H + FAD)] [40], mapped strictly within the somatic and axonal regions. We found that ORR increased significantly in day-10 animals within both cellular compartments, whereas day-5 animals remained statistically indistinguishable from the day-1 baseline, indicating a late-life transition toward a more oxidized neuronal microenvironment. Collectively, these results show that aging is accompanied by concurrent changes in neuronal mitochondrial states, morphology, and redox balance. The distinct responses observed in the soma and axon reveal compartment-specific mitochondrial aging within individual neurons, highlighting the capacity of our 2p-FLIM workflow to resolve *in vivo* subcellular mitochondrial remodeling.

## 3 Discussion

Here, we established a robust *in vivo* framework that integrates TMRM-based 2p-FLIM, MitoSLIT segmentation, phasor-intensity clustering, and spatial mapping to resolve mitochondrial functional states within living, intact *C. elegans*. This integrated approach distinguishes mitochondrial states and spatial heterogeneity that cannot be easily separated by fluorescence intensity alone. Furthermore, we identified sample-mounting-induced hypoxia as a critical confounding variable that rapidly alters mitochondrial morphology and bioenergetic readouts, underscoring the necessity of strict microenvironmental control during live imaging. Under these controlled conditions, we demonstrate that mitochondrial populations are highly divergent across different tissues in young adults and do not converge on a uniform bioenergetic end-point during aging. Instead, they follow distinct spatiotemporal trajectories across somatic tissues and even within the discrete subcellular compartments of individual neurons.

To address the challenge of *in vivo* characterizing mitochondrial heterogeneity across diverse tissues and within individual neurons at an organismal level, we leveraged the advantages of two-photon fluorescence lifetime imaging microscopy (2p-FLIM) combined with the commercially available mitochondrial dye TMRM. Compared to other widely used cationic potentiometric dyes such as TMRE and Rhodamine 123, TMRM exhibits the lowest non-specific membrane binding and minimal inhibition of cellular respiration [24], making it best suited for physiological imaging. Fluorescence lifetime provides a complementary readout sensitive to the local molecular environment of the fluorophore [18], adding additional resolving power to characterize mitochondrial states. Although novel mitochondrial probes [14–16] and FLIM probes [41–43] continue to be developed, most are not commercially available and have been validated primarily in cultured cells. Our framework instead uses routinely available mitochondrial dye TMRM to simultaneously measure mitochondrial architecture and spatial organization, ΔΨ*m*-associated fluorescence intensity, and lifetime-defined physicochemical state in intact *C. elegans*. We validate the high organelle specificity of TMRM in living *C. elegans*, demonstrating strong colocalization with mito::GFP from somatic tissues to single neurons. This integrated analysis identified eight data-driven phasor-intensity states and revealed tissue– and compartment-specific patterns of mitochondrial remodeling that were not apparent from conventional intensity measurements.

Our findings emphasize that oxygen availability is not merely a technical variable during live *C. elegans* imaging, but a critical determinant of the baseline mitochondrial phenotype. Utilizing 2p-FLIM, we demonstrated that the transient hypoxia induced by glass-based immobilization methods is sufficient to alter both mitochondrial spatial architecture and bioenergetic readouts in approximately 30 minutes. Gas-permeable coverslips largely prevented these changes, supporting restricted gas exchange as their primary cause. Thus, sample-mounting-induced hypoxia is a critical experimental confounder because it can produce acute mitochondrial-state transitions that may be misinterpreted as baseline physiology, aging, or experimentally induced stress.

Beyond this methodological implication, the observed hypoxia response provides insight into how mitochondria remodel under physiological and pathological oxygen limitation. Prior studies in *C. elegans* similarly showed that oxygen deprivation induces DRP-1-dependent mitochondrial fission in neurons [44] and disrupts mitochondrial morphology, ΔΨ*m*, and proteostasis [45]. Studies in mammalian systems also show that hypoxia and ischemia induce mitochondrial fragmentation, abnormal enlargement, and ΔΨ*m* loss, suggesting conserved physiological responses of mitochondria to oxygen limitation [46, 47]. Hypoxia-related signaling is also linked to aging. HIF-1 regulates lifespan and stress resistance in *C. elegans* in a context-dependent manner [48, 49], while age-associated NAD+ decline induces a pseudohypoxic state that disrupts nuclear–mitochondrial communication in mammals [50]. Consistent with these connections, acute hypoxia and aging produced partially overlapping mitochondrial signatures in our analyses, including reduced TMRM intensity, altered fluorescence lifetime, and more punctate mitochondrial organization. This phenotypic overlap raises the possibility that mitochondrial remodeling pathways activated by oxygen limitation also contribute to age-associated mitochondrial changes. Combining this imaging framework with genetic manipulation of oxygen-sensing and hypoxia-response pathways should help test this possibility.

Under the gas-permeable imaging conditions, calibrated TMRM intensity declined with age in the gonad, hypodermis, and pharynx, but fluorescence lifetime, state composition, and spatial organization followed distinct trajectories. Gonadal state redistribution was comparatively uniform. In the hypodermis, day-5 animals showed diffuse redistribution, whereas day-10 animals exhibited more pronounced subregional enrichment. In the pharynx, distinct states emerged within internal regions of the terminal bulb relative to the surrounding muscle. Thus, a shared decline in TMRM intensity accompanied fundamentally different forms of mitochondrial remodeling. These findings extend growing evidence that aging proceeds asynchronously across tissues to the level of mitochondrial physiology [8, 51–53].

Shifts in TMRM fluorescence lifetimes directly report mitochondrial microenvironmental alterations. Under high ΔΨ*m*, TMRM accumulates in the mitochondrial matrix, exhibiting brighter fluorescence signals, where FLIM resolves a subtle concentration-induced lifetime shortening driven by local quenching. Conversely, a transition toward longer lifetimes indicates TMRM redistribution from this partially quenched matrix environment into less polar, unquenched cellular fractions, a phenomenon characteristic of ΔΨ*m* collapse or mitochondrial dysfunction. This prolongation of fluorescence lifetime and concurrent decrease in calibrated intensity following mitochondrial depolarization is reported in the literature using cell lines [54] and consistent with our *C. elegans* data following treatment with the potent mitochondrial uncoupler FCCP (Fig. S11A). Accordingly, our *in vivo* aging data reveal a progressive increase in TMRM lifetimes paired with a concomitant decrease in calibrated intensity across all examined somatic tissues. However, post-reproductive gonads in day-5 and day-10 animals deviate from this systemic trend. In senescent gonads undergoing programmed reproductive death, compromised mitochondria retain TMRM and may become sequestered within autophagic and lysosomal compartments [34]. Within this distinct, hydrophobic, and degradative microenvironment, localized TMRM molecules display a shortened lifetime that, crucially, occurs independently of matrix self-quenching, as evidenced by the results that FCCP treatment did not induce lifetime changes in the mitochondrial region of the gonads undergoing self-destructive reproductive death (Fig. S11B).

Because mitochondria act both as a driver and as a readout of aging, these tissue-specific trajectories can be compared with previously described patterns of tissue decline in *C. elegans*. The early gonadal shift coincides with advanced reproductive aging, when oocyte quality declines and germline architecture is extensively remodeled [27, 32]. The subregional pattern in the hypodermis parallels ultrastructural reports that the aging hypodermis thins and loses integrity locally, with membrane degeneration, lipid accumulation, and cytoskeletal deterioration [51]. The pharyngeal pattern is consistent with the reported structural decline of the aging pharynx, including sarcopenia-like changes [28]. Tissue asynchrony has likewise been reported at the transcriptional level, where individual tissues display distinct aging trajectories and distinct sets of longevity regulators [53], consistent with the view that aging proceeds with distinct temporal profiles across tissues [8]. Our measurements extend these observations to organelle physiology.

Comparisons with environmental stress and defective mitochondrial dynamics further supported the tissue dependence of mitochondrial aging. Our results indicate that stressors and fission–fusion mutations did not reproduce a single aged state across tissues. In the high-energy-demand gonad, nearly all experimental perturbations drive mitochondrial phenotypes away from the day-1 baseline, with starvation inducing a profile that most closely recapitulates the post-reproductive, late-aging stage. Conversely, within the hypodermis, mitochondrial fission-fusion mutations mimic mid-stage aging, whereas late-stage aging more closely resembles acute stress conditions. Meanwhile, the pharynx displays a progressive, overlapping trajectory that shares features with both fusion-deficient states and oxidative stress conditions. To robustly validate these phenotypic relationships, the tissue-specific similarity heat maps shown in Fig. 5B were recomputed using alternative distance metrics, including the Hellinger distance [55] and the Aitchison distance [56]. Mantel tests [57] comparing these alternative metrics against the baseline Jensen-Shannon divergence framework (Fig. 5B) revealed uniformly high correlations, confirming that the emerging similarity structures reflect intrinsic data topology rather than distance-metric artifacts (Fig. S10).

The spatial organization of mitochondrial aging extended from tissue differences to subcellular compartments within individual neurons. Neuronal mitochondria differ in morphology and molecular composition between somatodendritic and axonal compartments [58–60]. Consistent with this compartmentalization, aging produced distinct changes in mitochondrial state and morphology between the soma and axon of AIY neurons. TMRM intensity declined in both compartments, but mitochondrial state distributions differed, with greater lifetime heterogeneity in the axon, while somatic mitochondria became shorter, rounder, and more compact. The optical redox ratio also increased in both compartments by day 10. These findings extend previous observations of compartment-specific mitochondrial aging in the mouse brain to a genetically identified neuron in a living animal, demonstrating that mitochondrial aging is shaped by the local subcellular environment [60].

In conclusion, the multi-layered descriptors extracted from 2p-FLIM data enable the identification of mitochondrial states in a spatiotemporal manner across diverse tissues, offering a powerful, *in vivo* framework for deep mitochondrial characterization in aging, neuroscience, cell metabolism, and development.

## 4 Methods

### 4.1 Strains

The *C. elegans* strains used in this study included wild-type Bristol N2, SJ4103 *zcIs14 [myo-3p::GFP(mit)]*, BXN723 *fzo-1(cjn20) II*, CU6372 *drp-1(tm1108) IV*, DA631 *eat-3(ad426) II; him-8(e1489) IV*, Han005 *sbsIs12[Pttx-3::rab-3::mCherry; Pttx-2::mitoGFP; ccGFP]*, and KQ2691 *nhr-91(tm4713) X; Ex[nhr-91p::nhr-91(genomic)::SL2::GFP, tdc-1p::mCherry, nmr-1p::mCherry]*. KQ2691 was kindly provided by the Kaveh Ashrafi laboratory at the University of California, San Francisco. Bristol N2, SJ4103, BXN723, CU6372, and DA631 were obtained from the Caenorhabditis Genetics Center (CGC), University of Minnesota. Nematodes used for measurements were synchronized using a two-generation egg-laying method. All synchronized animals were maintained at 20 ^◦^C on nematode growth medium (NGM) plates seeded with *Escherichia coli* OP50.

### 4.2 TMRM vital staining and experimental perturbations

Tetramethylrhodamine methyl ester perchlorate (TMRM; Invitrogen, T668) was dissolved in dimethyl sulfoxide (DMSO) to prepare a 200 *µ*M stock solution. TMRM staining plates were prepared one day before imaging by mixing freshly cultured *Escherichia coli* OP50 with TMRM and spreading the mixture onto nematode growth medium (NGM) plates at a final TMRM concentration of 200 nM. Plates were maintained overnight at room temperature in the dark. Unless otherwise specified, synchronized animals were stained for ∼1.5 h at 20 ^◦^C. For neuronal imaging, animals were stained for 3 h under the same TMRM concentration.

For FCCP treatment, animals were first incubated for 30 min on plates containing 0.1% DMSO and 200 nM TMRM, and then transferred to plates containing 20 *µ*M FCCP and 200 nM TMRM for an additional 1 h. Vehicle-control animals were incubated on plates containing 0.1% DMSO and 200 nM TMRM for 1.5 h. For oxidative stress conditions, animals were exposed overnight to paraquat dichloride hydrate (Sigma-Aldrich, 36541) at either 1 mM or 8 mg/mL (∼31 mM), followed by staining with 200 nM TMRM for ∼1.5 h. For heat stress, animals were incubated on plates containing 200 nM TMRM at 35 ^◦^ C for 2 h. For starvation, animals were incubated for 2 h in bacteria-free M9 buffer containing 400 nM TMRM. Mitochondrial-dynamics mutants were stained using the standard plate-based protocol.

Before imaging, animals were briefly transferred to unseeded NGM plates and then mounted under gas-permeable #1.5 polymer coverslips (ibidi) for live imaging. Animals were immobilized with 1 mM levamisole, except for aged and stress-treated animals, for which 0.25 mM levamisole was used.

### 4.3 Two-photon fluorescence lifetime imaging microscopy

The two-photon fluorescence lifetime imaging microscopy (2p-FLIM) measurements were performed on a Bruker Ultima Investigator microscope (Bruker, Billerica, MA) with two Hamamatsu H10770 Photomultiplier tubes (Hamamatsu Photonics, Shizuoka, Japan) for detection and a time-correlated single-photon counting (TCSPC) card (SPC-150 card, Becker & Hickl, GmbH, Berlin, Germany) for FLIM data acquisition. A 100-fs pulse laser with an 80 MHz repetition rate (Discovery Chameleon, Coherent, Santa Clara, CA) was used as the two-photon fluorescence excitation source. The excitation wavelength for TMRM and GFP was set at 850 nm for mitochondrial detection. An oil-immersion 60×/1.40 NA microscope objective (Nikon, Tokyo, Japan) was used for imaging. Emitted fluorescence was spectrally separated using a 565-nm long-pass dichroic mirror (Chroma Technology Corporation, Bellows Falls, VT). TMRM fluorescence was collected in the longer-wavelength detection channel through a 700-nm short-pass filter, whereas GFP fluorescence was collected through a 515/10-nm bandpass filter (Thorlabs, Newton, NJ). For imaging of the KQ2691 strain, the TMRM detection channel was additionally equipped with a 575-nm long-pass emission filter (Thorlabs, Newton, NJ). Excitation power was adjusted as needed to achieve sufficient photon statistics for FLIM analysis. The maximum detector constant-fraction discriminator (CFD) count rate was maintained at approximately 1×10^6^ counts s^-1^ to minimize detector saturation and TCSPC pile-up.

### 4.4 Calibrated TMRM intensity

Because the autofluorescence is ubiquitous across tissues and spans a broad emission range (∼400–600 nm)[25], which is largely overlapping with the TMRM emission window (∼570–620 nm), the best two-photon excitation wavelength of TMRM that yields the highest intensity ratio between TMRM and *C. elegans* intrinsic autofluorescence was determined experimentally. The excitation wavelength of the 100-fs tunable Ti:Sapphire laser was scanned from 790 to 1010 nm with a 20-nm increment. For each increment, the two-photon excitation fluorescence (TPEF) images of the same 1-day adult wild-type worms with or without TMRM staining were recorded. Under identical laser power, detection settings, pixel dwell time, and with compensated wavelength-dependent Group Delay Dispersion (GDD) for the two-photon excitation, the 850nm wavelength provided the highest TMRM fluorescence intensity, with on average more than one order of magnitude stronger than autofluorescence between the living adult worms with and without TMRM staining (Fig. S2). This wavelength-dependent intensity profile also served as a reference to define an intensity threshold, where pixel intensity below this threshold was considered indistinguishable from autofluorescence and thus excluded from subsequent 2p-FLIM lifetime phasor analysis.

To minimize laser system fluctuations and to normalize the TMRM intensity under diverse experimental conditions, the TMRM intensity and lifetime were carefully calibrated with the standard reference, freshly prepared 100 *µ*M pH11 fluorescein aqueous solution [61, 62], for every 2p-FLIM measurement under the 850 nm excitation. The calibrated TMRM intensity was thus defined as the measured TMRM intensity normalized by the fluorescence intensity of freshly prepared fluorescein standard (100 *µ*M in pH 11 aqueous solution) that was measured in every 2p-FLIM experiment.

### 4.5 Confocal imaging and optical redox ratio (ORR) analysis

Animals were imaged using a Zeiss LSM 980 NLO laser-scanning confocal microscope equipped with a 63×/1.4 NA oil-immersion objective. Worms were immobilized with 1.1 mM levamisole and mounted on a gas-permeable #1.5 polymer coverslip (ibidi) for live confocal imaging. Four-channel confocal z-stacks were acquired to simultaneously identify the target neuron and measure endogenous metabolic autofluorescence. GFP and mCherry signals were used to define the mitochondria and neuronal region, while NAD(P)H and FAD autofluorescence were collected for optical redox analysis. NAD(P)H was excited at 405 nm and detected from 430–470 nm, and FAD was excited at 445 nm and detected from 560–650 nm.

For analysis, z-slices containing the target neuron were selected and sum-intensity-projected. A neuron-specific mask was generated from the overlapping GFP and mCherry signals and applied to the corresponding projected NAD(P)H and FAD images. Image processing and intensity quantification were performed in ImageJ/Fiji. Mean fluorescence intensities of the z-projected images within the masked region were measured, and the optical redox ratio (ORR) was calculated for each neuron as: ORR = FAD / [FAD + NAD(P)H], where FAD and NAD(P)H represent the mean fluorescence intensities within the same neuronal mask.

### 4.6 Mitochondria Segmentation with Locally Iterative Thresholding (MitoSLIT)

Mitochondria Segmentation with Locally Iterative Thresholding (MitoSLIT) is designed to detect TMRM-stained mitochondria through a multi-scale, probability-based segmentation framework. All computational procedures were implemented using Python (v.3.12), leveraging the scikit-image (v.0.24.0) [63] and SciPy (v.1.14.0) [64] libraries. The MitoSLIT pipeline proceeds as follows:

**1.** Preprocessing and Padding: To mitigate edge artifacts during sliding-window operations, the TMRM fluorescence intensity image undergoes reflective (mirror) edge padding (Fig. S4, Step 1).
**2.** Adaptive Parameterization: The image is partitioned into an ensemble of blocks across varying spatial scales (the yellow squares shown in Fig. S4, Step 2). A histogram of the mean pixel intensities per block is generated and modeled using a sigmoid function. To stabilize thresholding across high-intensity regions, this sigmoid is inverted to define a threshold plateau, establishing a non-linear mapping between local mean intensity and adaptive threshold values (Fig. S4, Step 3).
**3.** Iterative Local Binarization: A global threshold is first calculated using Otsu’s method, a popular and computationally efficient image thresholding method [29], to establish a baseline contrast. Subsequently, an iterative sliding window traverses the image (Fig. S4, Step 2). For each window position, a locally adaptive threshold is calculated as a function of the local mean intensity (Fig. S4, Step 3). Pixels exceeding this dynamic threshold are binarized as foreground (1), while those below are assigned to the background (0).
**4.** Probability Heatmap Generation: The binary outputs from all sliding window iterations and scales are spatially integrated (summed) to generate a probability heatmap (Fig. S4, Step 4). This map represents the statistical likelihood of mitochondrial occupancy based on local contrast rather than an absolute intensity. This approach ensures the detection of “dim” mitochondria with a low TMRM signal intensity that would otherwise be omitted by static global thresholding.
**5.** Final Segmentation: The probability heatmap is segmented using Maximum Entropy Thresholding [65] (Fig. S4, Step 5). The Maximum Entropy Thresholding algorithm identifies the optimal threshold by maximizing the inter-class entropy between foreground and background, providing a robust separation based on the information content of the probability distribution.

### 4.7 Mitochondrial morphological analysis

#### 4.7.1 Fractal dimension analysis

Box-counting fractal dimension was calculated from MitoSLIT-segmented mitochondrial binary masks to quantify the global space-filling complexity of mitochondrial patterns [66]. Mitochondrial foreground pixels were first restricted to the corresponding tissue ROI, and pixels outside the ROI were excluded. The ROI-restricted binary mask was then tiled with square boxes of increasing size, and the number of partially occupied boxes containing mitochondrial signal was counted at each scale. The fractal dimension, *D*0, was estimated as the negative slope of the linear fit between log(box count) and log(box size). For each tissue, a fixed box-size range was used across all ages and perturbation conditions. Gonad and pharynx were analyzed using box sizes of 4, 8, 16, and 32 pixels, whereas hypodermis was analyzed using box sizes of 4, 8, 16, 32, and 64 pixels. Fractal dimension values were compared within each tissue using Welch’s t-test and visualized using violin plots with internal box plots.

#### 4.7.2 Rotationally-invariant local binary pattern analysis

Local binary pattern (LBP) texture analysis [67] was performed on MitoSLIT-denoised mitochondrial fluorescence images to quantify scale-dependent mitochondrial texture patterns. For each image, the corresponding tissue ROI mask was used to restrict the analysis to the tissue region. Fluorescence intensities were rescaled based on each image’s intensity range and converted to 8-bit images before LBP calculation.

Rotationally invariant uniform LBP histograms were calculated at two spatial scales, with radii of 3 pixels for fine-scale texture and 10 pixels for coarse-scale texture. These radii corresponded to neighborhood diameters of approximately 1.2 and 4 *µ*m, respectively. The fine scale was selected to capture local intensity contrasts at about the scale of individual mitochondrial structures, whereas the coarse scale captured higher-order organization across multiple neighboring mitochondria. The same radii were applied to all tissues, ages, and perturbation conditions. For each radius, the number of sampling points was set to eight times the radius, and the resulting LBP codes were summarized as normalized frequency histograms. To avoid boundary artifacts, ROI masks were eroded by the corresponding LBP radius before feature extraction, so that only pixels whose full LBP neighborhood remained inside the tissue ROI were included. Image-radius combinations with fewer than 200 valid pixels after ROI erosion were excluded from that radius-specific analysis. Selected LBP bins representing spot-like, edge-like, ring-like, and non-uniform texture patterns were used for group-level comparison. LBP features were compared within each tissue using Welch’s t-test and visualized using selected-bin plots with individual image-level measurements.

#### 4.7.3 Neuronal mitochondria morphology analysis

Neuronal mitochondrial morphology was quantified from MitoSLIT-generated binary masks based on the GFP mitochondrial signal. Because neuronal regions were small and imaged at high magnification, morphology was analyzed at the level of individual segmented mitochondrial objects rather than using tissue-scale pattern analysis. Binary masks were connected-component labeled and size-filtered to remove small noise particles. For each segmented object, morphological descriptors were extracted, including area, convex area, filled area, equivalent diameter, extent, Euler number, perimeter, Crofton perimeter, solidity, major and minor axis lengths, orientation, and eccentricity. Derived metrics included aspect ratio, calculated as major axis length divided by minor axis length, and circularity, calculated as 4*π* × area / *perimeter*^2^. Young and aged groups were compared using Welch’s t-test. Significant morphology parameters were visualized as group means with standard deviation error bars.

#### 4.7.4 Computational analysis environment

Fractal dimension analysis, LBP texture analysis, and neuronal mitochondrial morphology analysis were performed using custom Python scripts. Analyses were conducted in Python 3.10.14 using NumPy 1.26.4 [68], pandas 2.2.3, SciPy 1.15.3 [64], scikit-image 0.25.2 [63], Matplotlib 3.7.3, and tqdm 4.67.1. Statistical comparisons were performed using Welch’s t-test implemented in SciPy.

### 4.8 Multiparametric similarity analysis

Similarity analyses were performed in Python (3.11) with NumPy (2.26) [68] and SciPy (1.15) [64], separately within each tissue across the ten aging, stress, and mutant conditions. For each image, two feature blocks were assembled: the mitochondrial-state cluster fractions (based on G, S, and I*calibr*) and morphology descriptors comprising the fractal dimension (*D*0) and LBP features. For the cluster-fraction block, the distance between two conditions was defined as the generalized energy distance [69] between their distributions of per-image vectors using the square-root Jensen– Shannon divergence as the base metric [70]. For the morphology block, features were z-scored, and an analogous energy distance was computed with a Euclidean base metric. The two condition×condition matrices were combined into a single distance matrix (Fig. 5B). For validation, the class-fraction block was recomputed using the Hellinger distance [55] and the Aitchison distance [56], and the resulting maps were compared to the Jensen–Shannon version by the Mantel test [57].

## Acknowledgements

This work was funded by NIH/NIA R21 (R21AG086974) to W-WC, SH, and MC, and by NIH/NIA R01 (R01AG081270) and Hevolution/AFAR New Investigator Award (AGR00030264) to SH. The authors thank the Kaveh Ashrafi laboratory at the University of California, San Francisco, for kindly providing the KQ2691 strain. The authors thank the microscopy core facility and Dr. Danielle Scheff and Dr. Sandy Hsieh at the Parker H. Petit Institute for Bioengineering and Bioscience at the Georgia Institute of Technology for providing access to and assistance with shared microscopy equipment.

## Author contributions

W-WC, WT, SH, and MC conceived and designed the study. WT and W-WC conducted 2p-FLIM measurements. WT conducted ORR experiments and analysis, and neuronal mitochondrial morphology analysis. HX conducted the similarity analysis. WT and SB performed *C. elegans* culture and material preparation. WT, HX, and W-WC analyzed the 2p-FLIM data, with discussions and contributions from SH, MC, and SB. WT, HX, W-WC, SH, and MC wrote the manuscript. All authors reviewed the manuscript.

