## Supplemental figures for "Multiparametric *in vivo* mapping reveals tissue-specific mitochondrial aging trajectories"

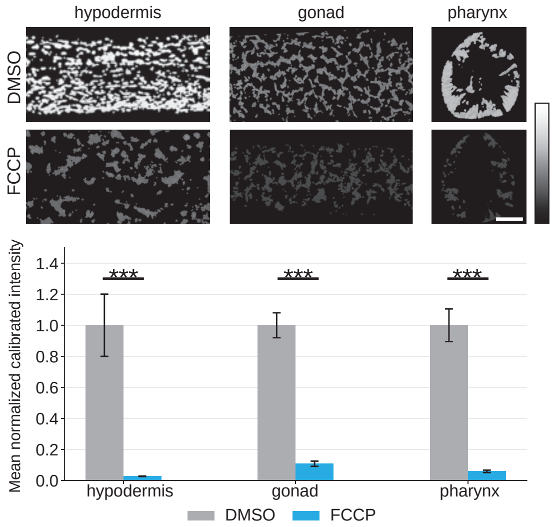


**Fig. S1**: **FCCP-induced mitochondrial depolarization reduces TMRM fluorescence intensity across tissues** Representative TMRM images (top) and quantification of calibrated fluorescence intensity (bottom) in the hypodermis, gonad, and pharynx of worms treated with 0.1% DMSO (vehicle) or 20 $\mu$M FCCP. Intensities were calibrated to laser power and normalized to the mean of the corresponding DMSO group. Data are mean $\pm$ SEM; DMSO, $n = 10$ worms per tissue; FCCP, $n = 8$, 9, and 8 worms for the hypodermis, gonad and pharynx, respectively. Two-sided Welch's t-test. ns, not significant; $p < 0.05$ (*), $p < 0.01$ (**), and $p < 0.001$ (***). Scale bar = 10 $\mu$m.

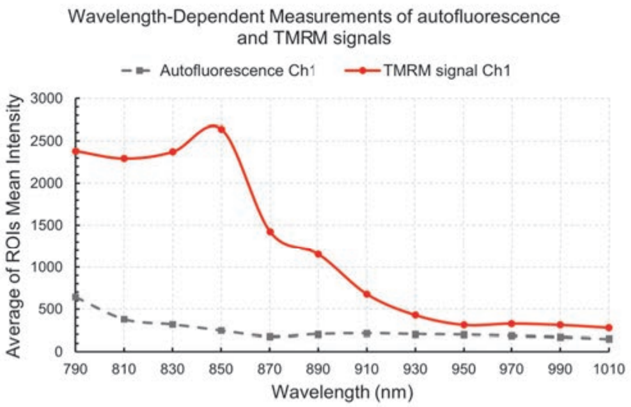


**Fig. S2**: **Wavelength-dependent two-photon excitation of TMRM** Average ROI fluorescence intensity was measured in 1-day-old adult wild-type animals with or without TMRM staining across excitation wavelengths from 790 to 1010 nm. At 850 nm, the TMRM signal is, on average, more than one order of magnitude stronger than autofluorescence under identical laser power, detection settings, pixel dwell time, and with compensated wavelength-dependent Group Delay Dispersion (GDD) for the two-photon excitation, indicating a minor contribution of the intrinsic non-intestinal autofluorescence to the measured TMRM signal.

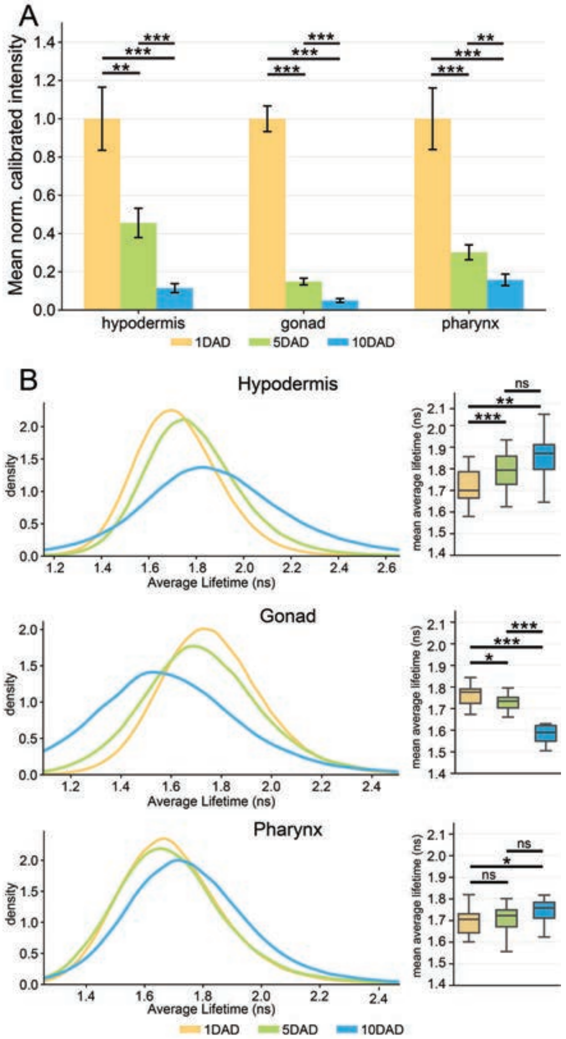


**Fig. S3**: **Quantitative analysis of tissue-dependent TMRM fluorescence intensity and lifetime changes during aging** (A) Mean TMRM fluorescence intensity was quantified from ROI-masked mitochondrial regions in the hypodermis, gonad, and pharynx of day-1, day-5, and day-10 adult *C. elegans*. Raw fluorescence intensity was calibrated by laser power using fluorescein-based calibration and normalized to the day-1 mean within each tissue. Bars indicate mean ± SEM. (B) Average fluorescence lifetime distributions were analyzed from ROI-masked mitochondrial pixels for each tissue and age group. Density plots show pooled pixel-level average lifetime distributions, while boxplots summarize the image-level mean average lifetime. Sample sizes at day 1, day 5, and day 10 were $n = 31, 25$, and 14 images for hypodermis, $n = 19, 16$, and 6 for gonad, and $n = 24, 16$, and 21 for pharynx, respectively. Statistical comparisons were performed using Welch's t-test. ns, not significant; $p < 0.05$ (*), $p < 0.01$ (**), and $p < 0.001$ (***).

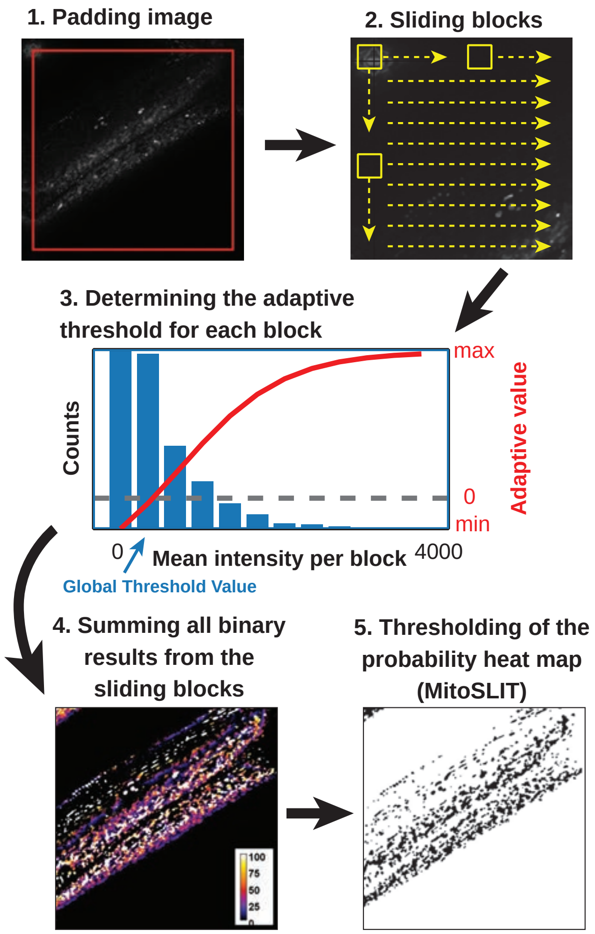


**Fig. S4: Flowchart of Mitochondria Segmentation with Locally Iterative Thresholding (MitoSLIT)** Image processing steps for detecting TMRM-labeled mitochondria with high and low intensities in an image.

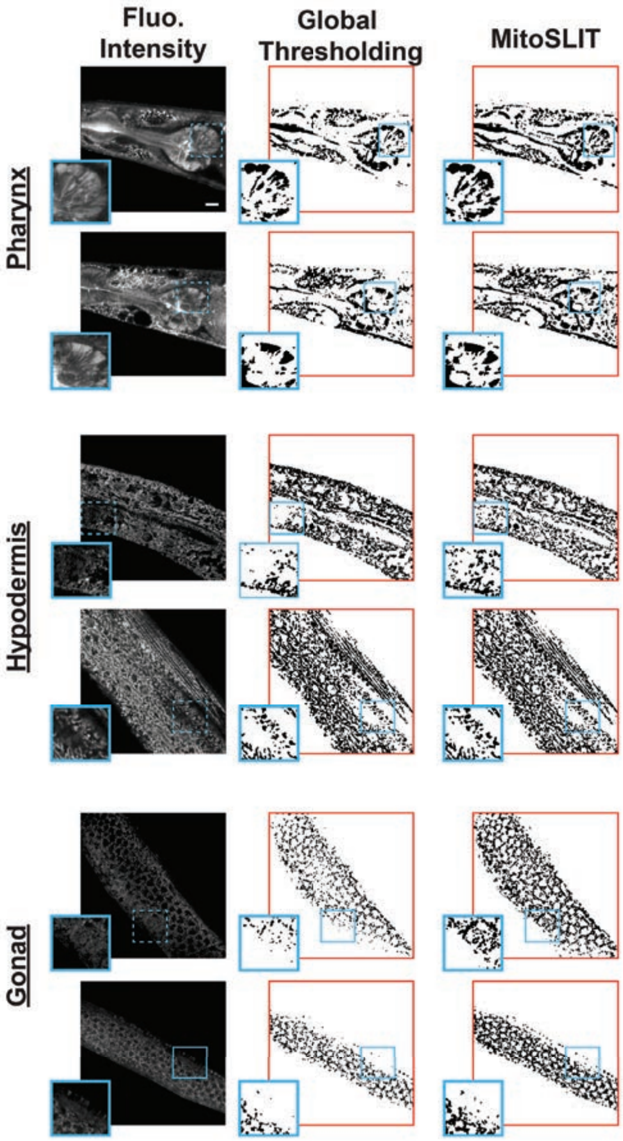


**Fig. S5**: **Results of applying global thresholding and MitoSLIT to images of day-5 wild-type worms with *in vivo* TMRM staining** Compared to the global thresholding approach, MitoSLIT can detect both strong and weak signals in the same images. Scale bar = 10 $\mu$m.

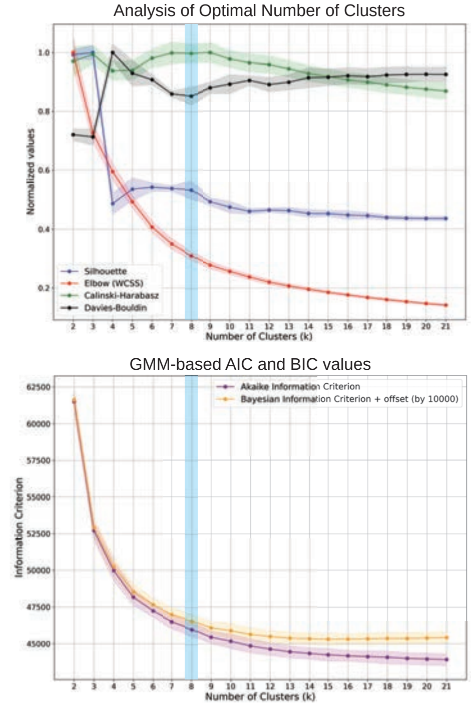


**Fig. S6**: **Analysis for optimal number of clusters** Normalized silhouette, within-cluster sum of squares (WCSS), Calinski–Harabasz, and Davies–Bouldin scores obtained using K-means clustering across k=2–21 (upper panel).Akaike information criterion (AIC) and Bayesian information criterion (BIC) values obtained using Gaussian mixture models (lower panel). Curves show the mean of 30 random subsampling iterations, with shaded regions indicating ± SD. The blue-shaded region marks the selected solution with k=8, which was broadly supported across clustering and model-selection criteria.

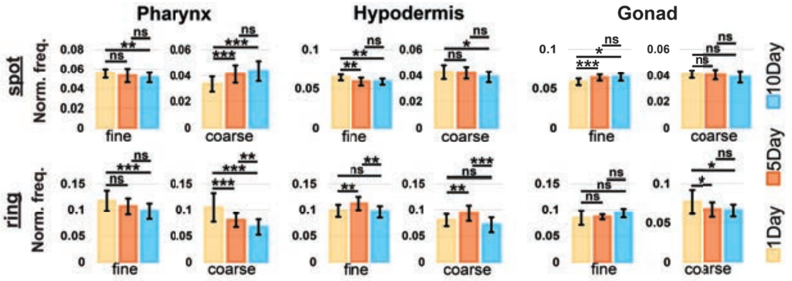


**Fig. S7**: **LBP analysis of aging tissues** Local binary pattern analysis of mitochondrial texture features at fine and coarse spatial scales. Bar plots summarize normalized frequencies of spot-like and ring-like texture features. Bars indicate mean ± SD. Pairwise comparisons were performed using two-tailed Welch's t-test. ns, not significant; $p < 0.05$ (*), $p < 0.01$ (**), and $p < 0.001$ (***).

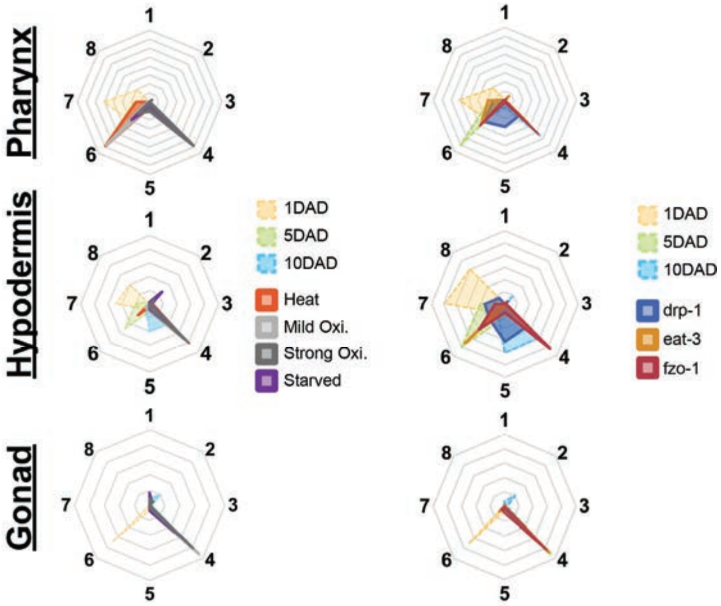


**Fig. S8**: **Mitochondrial population distribution of mutant and stress conditions** Radar plots show the relative abundance of the eight mitochondrial clusters in the pharynx, hypodermis, and gonad. Aging groups (1DAD, 5DAD, and 10DAD for 1-day, 5-day, and 10-day adults, respectively) are compared with acute stress conditions (left) and mitochondrial-associated mutants (right). Each axis represents one cluster, and radial distance indicates its relative abundance.

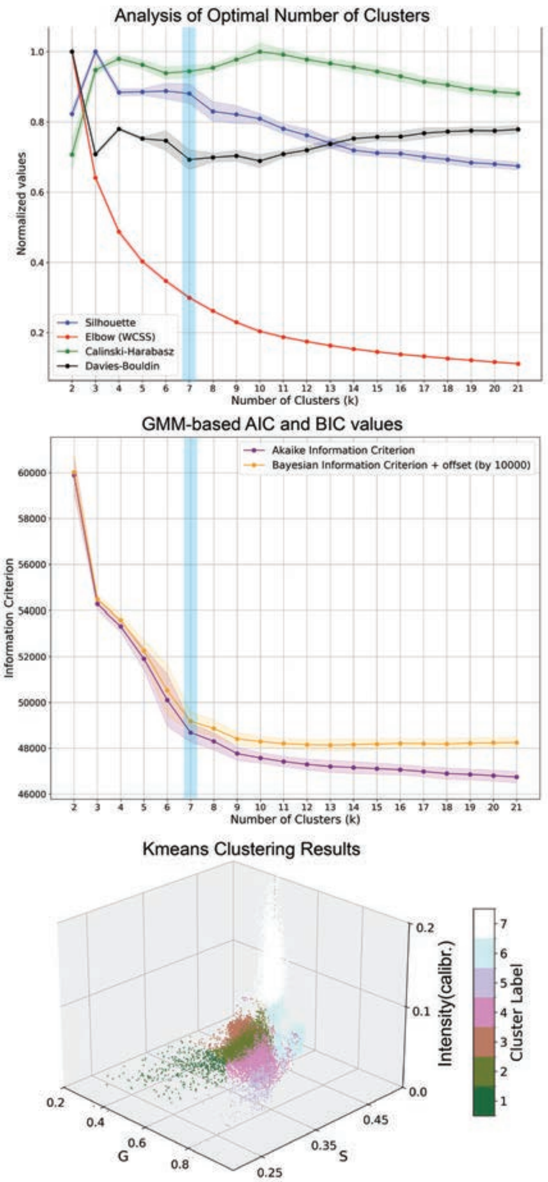


**Fig. S9**: **Cluster analysis of neuronal mitochondria** Normalized silhouette, within-cluster sum of squares (WCSS), Calinski–Harabasz, and Davies–Bouldin scores across different values of k (upper panel). AIC and BIC values obtained using Gaussian mixture models. Curves show the mean across 30 random subsampling iterations, with shaded regions indicating $\pm$ SD (middle panel). Overall, the clustering evaluation metrics and model-selection criteria supported k=7, as indicated by the blue-shaded region. Three-dimensional visualization of the seven K-means clusters based on the phasor coordinates G and S and calibrated fluorescence intensity (lower panel).

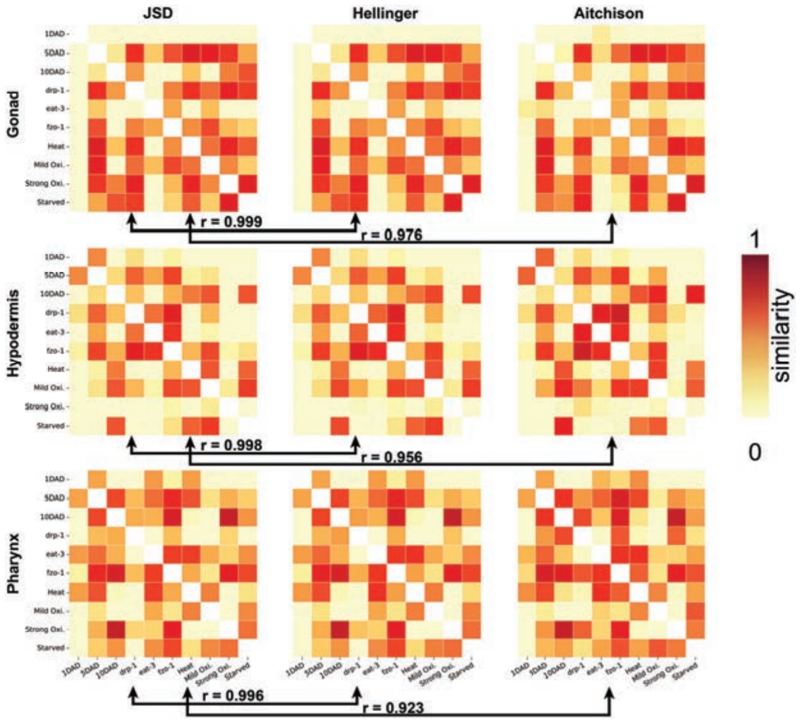


**Fig. S10: Similarity maps are robust to the choice of distance metric.** Similarity score maps for each tissue (rows: gonad, hypodermis, pharynx) computed with three different compositional distances on the mitochondrial class-fraction block (columns: Jensen-Shannon Divergence, Hellinger, Aitchison). For each alternative metric, the Spearman rank correlation is reported uniformly high between the alternatives, indicating that the similarity is a property of the data rather than an artifact of the chosen distance.

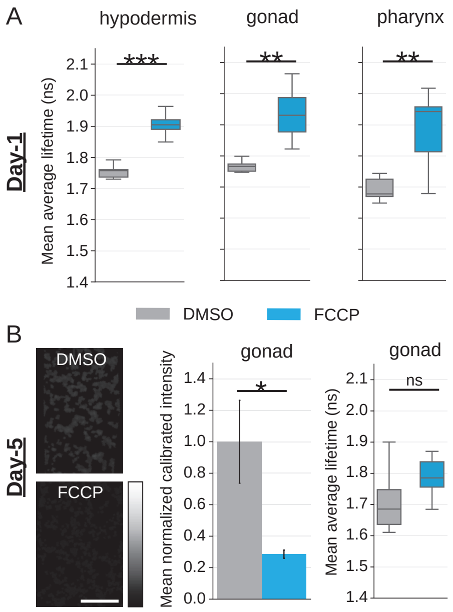


**Fig. S11**: **FCCP-induced changes in TMRM fluorescence lifetime and intensity.** (A) Mean TMRM fluorescence lifetime in the hypodermis, gonad and pharynx of day-1 adult worms treated with 0.1% DMSO or 20 $\mu$M FCCP. For DMSO, $n = 10$ worms per tissue; for FCCP, $n = 10$, 9, and 8 worms for the hypodermis, gonad, and pharynx, respectively. (B) Representative calibrated TMRM intensity images (left) and quantification of normalized calibrated intensity (middle) and fluorescence lifetime (right) in day-5 adult gonads under the same treatment conditions ($n = 8$ worms per condition). Intensity was normalized to the mean DMSO value. Data are mean $\pm$ SEM. Two-sided Welch's t-test. ns, not significant; $p < 0.05$ (*), $p < 0.01$ (**), and $p < 0.001$ (***). Scale bar = 10 $\mu$m.
